# Learning a PRECISE language for small-molecule binding

**DOI:** 10.64898/2026.01.04.697581

**Authors:** Mert Erden, Xuting Zhang, Kapil Devkota, Rohit Singh, Lenore Cowen

**Author notes:** These authors contributed equally (Mert Erden, Xuting Zhang, Kapil Devkota).

## Abstract

Virtual screening of billion-scale compound libraries has become feasible through machine learning approaches. In particular, CoNCISE (RECOMB 2025) introduced drug quantization via codebooks, achieving highly scalable and accurate binary predictions. However, drug discovery requires understanding not just whether molecules bind, but where they bind and how to target specific sites. Here, we present PRECISE which leverages CoNCISE’s quantized small-molecule representations while operating on the target’s 3D structure as its input. The key innovation of PRECISE is reimagining drug-target interaction as compatibility between quantized drug embeddings and a latent representation of the target’s surface mesh, enriched with electrostatic and geometric features. PRECISE designs a novel surface representation, interpreted through a geometric deep learning architecture, enabling it to identify binding sites more accurately than state-of-the-art methods (DiffDock-L, Chai, and Boltz-2) while the codebook ensures billion-scale screening capability. Our formulation unlocks zero-shot generalization to complex targets such as metalloproteins and multi-chain complexes. To enable efficient integration with downstream docking workflows, we introduce Precise-MCTS, which combines fast Precise-based screening with selective Vina docking through an iterative Monte Carlo Tree Search approach. By providing both mechanistic understanding and massive scalability, PRECISE delivers capabilities that were previously mutually exclusive in virtual screening.

## 1 Introduction

In small-molecule based drug discovery, a foundational challenge is identifying strong binders to a protein target from the essentially infinite space of synthesizable compounds. Ultra-high-throughput virtual screening (uHTVS) offers a compelling solution by leveraging pre-enumerated libraries of commercially available molecules such as ZINC [20,21] (250 million), Enamine [16] (10.4 billion), and mCule [24] (140 million), where synthesizability and drug-likeness considerations are already addressed. The use of machine learning-derived protein and drug representations has enabled substantial advances in screening efficiency and accuracy [44,17]. We previously presented CoNCISE [13], which can screen billions of compounds in minutes, providing ranked lists of potential binders with state-of-the-art accuracy. However, existing uHTVS methods (including CoNCISE) remain limited to binary predictions.

Specifically, existing uHTVS methods remain fundamentally limited in incorporating structural information: they are unable to tell where the molecule will bind, or limit the screening to molecules that bind to a specific pocket. This is because the current approaches featurize proteins and drugs in learned latent spaces, mapping representations directly to binding probability without revealing where molecules bind or which residues drive interaction. This limitation hampers drug design tasks such as developing selective inhibitors that target the regulatory but not the catalytic domain of a kinase.

A related but distinct challenge affects many virtual screening approaches: the inability to generalize beyond single-chain protein targets to handle complex molecular assemblies like metalloproteins, proteinprotein interfaces, or proteins with cofactors and post-translational modifications (PTMs). This limitation extends beyond uHTVS to modern machine learning (ML) based docking methods [30,1,45,51,19], which often fail when confronted with non-monomeric targets. Molecular dynamics simulations potentially handle such targets, but require hours or days per molecule, making them unsuitable for screening even modest libraries. These limitations leave important classes of therapeutically relevant targets computationally inaccessible.

We introduce Precise, a structure-based uHTVS approach that represents binding targets as solvent-excluded surface meshes enriched with physico-chemical and geometric properties. This surface-based abstraction captures some of the essential physics of molecular recognition—local physicochemical features and surface curvature—while remaining computationally efficient for billion-scale screening. Precise solves two problems: a) predict the likely binding site(s) on a target through a surface-wide assessment of ligand accessibility, and b) given a target surface and a specific site on it, perform site-specific screening of massive compound libraries. Precise uses a geometric deep learning architecture and builds on CoNCISE’s hierarchical drug codes to maintain computational efficiency at scale. By training surface representations on 3D structural data while leveraging drug codes derived from binary binding assays, Precise bridges the complementary DTI modalities of structural biology and binding-affinity assays.

Precise uses the protein 3D structure to compute the surface mesh, but then operates purely on surface representations for the target, requiring no sequence information once the surface mesh is characterized. This enables direct zero-shot generalization to complex targets: a metalloprotein is simply a surface with altered electrostatic patterns from metal coordination, while a protein-protein interface in a ternary complex presents as an extended surface geometry spanning multiple chains. To our knowledge, Precise is the first uHTVS method capable of handling such complex molecular assemblies.

Leveraging recent innovations in 3D equivariant architectures, our data-efficient surface abstraction enables Precise to outperform the ML-based docking method DiffDock-L [7,6] and match co-folding methods such as Chai [46] and Boltz-2 [39] for binding site prediction accuracy. Precise can perform ultra-fast drug screens. On a large database like ZincDB-250M, Precise is able to achieve a median Vina [12] score of −8.8 across 65 targets, improving upon recent generative approaches like Rag2Mol [54] (median Vina score = −8.6). Critically, Precise is 12x faster and reports commercially available molecules while Rag2Mol reports molecules that need custom synthesis (i.e., not available for immediate purchase).

Our results suggest that Precise’s mechanism-aware screening at scale opens new avenues for drug discovery, democratizing access to labs without synthetic chemistry capabilities. Beyond traditional singleprotein targets, Precise unlocks new discovery modes, such as finding ligands that bind differentially to allosteric versus orthosteric sites or more strongly to disease-associated mutants. Our framework naturally extends to handle post-translational modifications, covalent inhibitors, and other complex scenarios that share the common thread of being representable as surfaces with defined properties.

## 2 Problem Formulation and Related Work

Ultra-high-throughput virtual screening (uHTVS) can be formulated in two distinct settings. The first takes a target protein and identifies potential small-molecule binders from a database of candidate ligands [38,37,28,18,45]. The second setting narrows this task: given both a target protein and a specific binding pocket, identify small molecules that bind at that location [47,12,14,32,53]. The two settings imply some critical tradeoffs: protein-level screening (Setting 1) can typically be solved just with sequence inputs, and often much faster than pocket-specific screening (Setting 2), but provides no mechanistic guarantee that the shortlisted binders will target the desired pocket.

Two methods presented at RECOMB 2025 illustrate this tradeoff. CoNCISE [13] (Fig 1a) addressed Setting 1 by discretizing the ligand space into 32, 768 (= 32×32×32) hierarchical codes. This quantization reduced screening to a database lookup problem, enabling searches through billions of compounds in minutes. CoNCISE achieved state-of-the-art accuracy for binary prediction of binding. Rag2Mol [54] tackled Setting 2 through structure-based generative modeling. The method retrieved ligands with CoNPLex [44] from compounded databases to discover scaffolds. Using high scoring scaffolds, they generated *de novo* ligands that targeted specific protein regions. While Rag2Mol achieved high pocket specificity through its generative approach, the approach does not extend to multimers, the initial search costs scale linearly with database size, and the final generated candidates may still not be synthesizable. These methods expose a fundamental tension in current uHTVS: scalability and mechanistic specificity have seemed mutually exclusive.

**Fig. 1:**
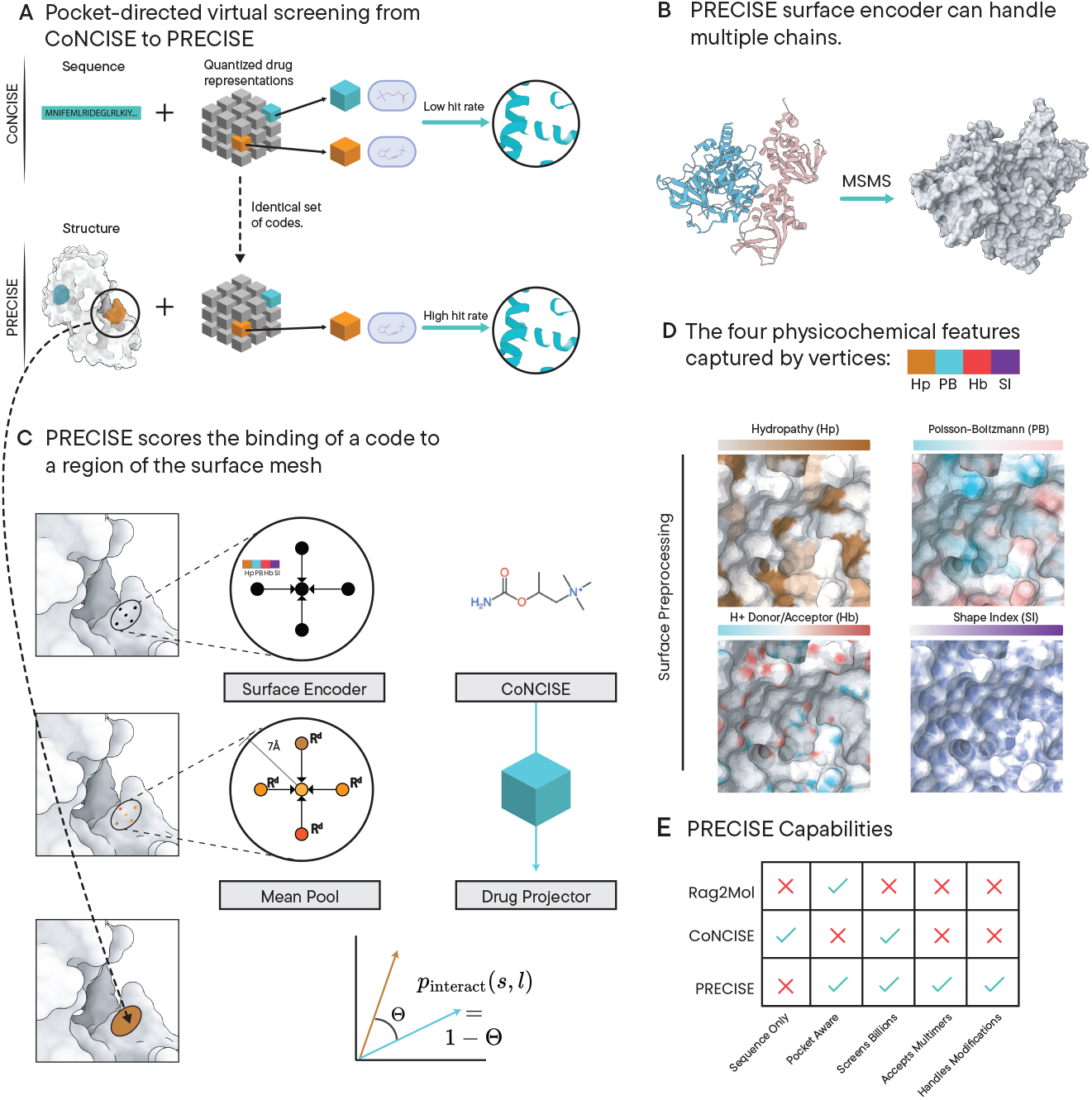
**Overview** A) Precise uses the same codebook index for small molecules as CoNCISE to enable ultra-fast screening of large databases but Precise takes as input a target’s 3D structure, rather than the protein sequence B) Precise’s surface-based approach allows incorporation of higher-order quarternary structures (in this case the target consists of two protein chains in a complex) without any model reconfiguration. C) The basic Precise operation computes the local score of a molecular code’s binding propensity to a surface vertex. These per-vertex scores are aggregated and the molecule is predicted to bind to the specified pocket if many of the sampled vertices in the patch have high scores against the molecule. D) Precise’s surface representation consists of four physico-chemical and geometric properties. Once these are computed, we don’t further utilize the protein sequence. E) Precise relieves the tension between scalability and targeted ligand screening, a critical gap in uHTVS, while also generalizing to composite surfaces.

Precise resolves this tension by combining CoNCISE’s quantized ligand representation with a surface-based protein representation (enriched with local physicochemical descriptors) that captures pocket-level binding determinants. This enables directed screening at user-specified locations on the target. Crucially, representing the target using *only* surface-based features means Precise generalizes automatically to handle additional structural augmentations such as oligomerization state, metal ion presence, or post-translational modifications, which other methods struggle with. With a surface abstraction that generalizes to these complex targets, Precise addresses a critical gap in current drug design methodologies.

## 3 Methodology

The core Precise model performs a single elementary operation: predicting whether a small molecule binds at a specific protein surface patch. This prediction integrates two representations. First, small molecules are indexed using the DTI-binding aware hierarchical codes learned by CoNCISE (Figure 1A and Section 3.2). These codes co-embed drugs with protein language model-based protein representations, yielding drug encodings that capture binding-relevant molecular properties. Notably, code construction in CoNCISE relied only on protein sequence information. Nonetheless, we hypothesize that the codes will generalize well to the structure-based setting. Second, the target surface is extracted from its 3D structure and characterized through local geometric and physicochemical properties (Figure 1B-D and Section 3.1). For each surface vertex, we compute both the local surface geometry and physicochemical features: hydropathy, electrostatics, hydrogen bonding potential, and shape index.

Precise learns to predict binding by training on structures of protein-ligand complexes. We work only with the bound conformation; we do not model conformational changes between apo and holo states. Following previous co-embedding approaches like ConPLex [44] and CoNCISE [13], which mapped molecular fingerprints and protein sequences into a shared embedding space, Precise co-embeds drug codes and surface patches to predict binding. We define binding as proximity within 7 Å of a surface vertex, following the patch radius recommended by Gainza et al. [15].

These hyper-local predictions generate a binding propensity “heat-map” across the protein surface. Scattered predictions indicate no coherent binding site, while spatially aggregated high-scoring vertices define a binding patch.^4^ Our aggregation procedure enables pocket prediction (Section 3.4, Figure 2). Beyond pocket-level predictions, the maximum pocket score also provides a whole-molecule DTI prediction. On the PLINDER benchmark [11], Precise achieves an area under precision-recall curve (AUPR) of 0.233 for binary DTI prediction, outperforming ConPLex (AUPR = 0.192) but trailing CoNCISE (AUPR = 0.472) (Section 3.6).

**Fig. 2:**
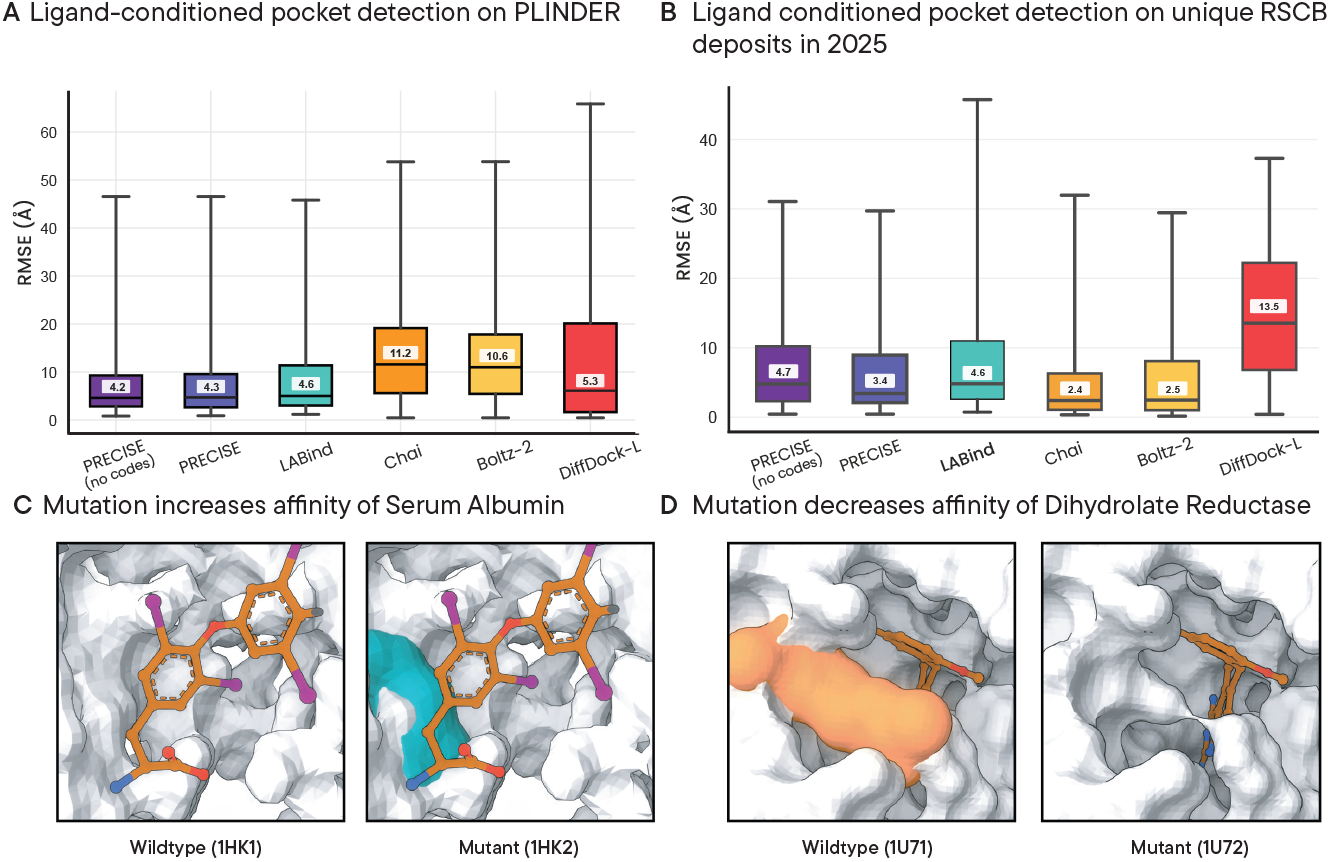
Ligand-driven capabilities of Precise. A) Precise is able predict binding pockets of structures on low sequence- and pocket-homology test examples. B) Under a time-based split Precise maintains similar performance. C) Precise detects the R218H mutation of human serum albumin. The mutation from arginine to histidine relaxes the sterics around the binding pocket [40] D) Mutation of leucine (in orange) to arginine in reductase loosens the binding pocket, lowering the binding affinity of the mutant. [5]

Precise offers the speed advantage of CoNCISE, combined with the pocket-oriented screening capability of methods like Rag2Mol [54] (Figure 1E). Moreoever, it is able to handle composite targets that neither sequence-based methods nor structure-based generative methods can address: multimeric assemblies, metalloproteins, and targets with post-translational modifications. These capabilities emerge naturally from the surface-based representation, requiring no model reconfiguration (Section 3.4).

### 3.1 Surface representation

Given a protein structure, its solvent-accessible surface mesh is generated using *Michael Sanner’s Molecular Surface (MSMS)* algorithm [43] and enriched with local physicochemical features (Figure 1 and Section A.2 in the Supplement). We hypothesize that a simple surface representation that captures local physicochemical and curvature properties is sufficient to characterize localized ligand specificity. We use MaSIF [31] to compute four surface features per mesh vertex: (a) Hydropathy as computed using the Kyte-Doolittle index [27], (b) Poisson-Boltzmann continuum electrostatics as computed by the APBS software [23], (c) hydrogen bonding characteristics [26] using RDKit, and (d) the local shape index [25]. Notably, for proteins in complex with other entities (e.g., metal ions, ligands, or even other proteins), Marchand et al. [31] recently described **MaSIF-neosurf** which extends these characterizations to such composite entities. By using this newly-developed featurization, our surface-ligand binding can generalize to such composites and even multi-chain complexes. This motivates our design decision to disregard sequence information once the surface mesh has been characterized, in contrast to the previous surface-based formulation of LABind [55]. **Surface processing with GotenNet**. Each surface mesh comprises vertex coordinates and triangulated edges, making it naturally suited for graph-attention models that support rotational equivariance and local message passing. A recent 3D graph-attention framework called GotenNet [2] showed that it can cheaply and effectively model the intricacies of a 3D graph. They achieved this by leveraging effective geometric tensor representations, not relying on computationally expensive irreducible representations or Clebsch-Gordan transforms [2]. We therefore adopted it as the core component of our surface module. The module accepts three inputs derived from the functionally enriched surface mesh:

1. A list of 3D vertex coordinates,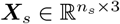;
2. A graph structure, *G*_*s*_ = (*V*_*s*_, *E*_*s*_), where |*V*_*s*_| = *n*_*s*_ defines the mesh connectivity; and
3. A set of per-vertex features, 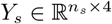, capturing localized physicochemical and geometric properties.

These three inputs (***X***_*s*_, *G*_*s*_, *Y*_*s*_) are provided to GotenNet, which performs equivariant message passing on the graph and captures both invariant and steerable (orientation-dependent) local properties (Supplement A.2). GotenNet returns two outputs: a) an equivariant representation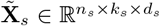, where *k*_*s*_ and *d*_*s*_ denote the steerable and invariant feature dimensions, respectively; and b) fully invariant embedding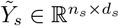.

To form the final invariant surface representation, the surface module normalizes the steerable component along the *k*_*s*_ dimension and concatenates it with the invariant output:

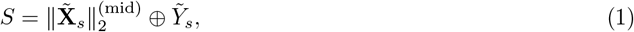

where 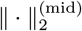 denotes the application of the *L*_2_-norm along the middle (*k*_*s*_) dimension, and ⊕ represents concatenation along the feature axis. After a non-linear down-projection from 2*d*_*s*_ to *d*_*s*_ dimensions, the resulting tensor 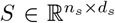 serves as the final invariant surface representation generated by the Precise surface module. A more detailed description of the surface block is provided in Supplement A.1.

### 3.2 Processing ligand SMILES

Given the ligand’s SMILES representation, Precise first computes its Morgan fingerprint (specifically, Extended-Connectivity Fingerprint ECFP-4 [42]) as input to its drug module. These fingerprints are then discretized using CoNCISE ‘s pretrained drug encoder. CoNCISE ‘s encoder was originally trained on binary DTI data derived solely from sequence–SMILES pairs, without any explicit structural information. Using finite scalar quantization [33], CoNCISE organized the entire space of small molecules into a 3-tier hierarchy, with 32 categories at each level, resulting in 32,768 (=32^3^) drug codes. Leveraging such a sequence-informed ligand representation for a structure-based surface prediction task is not straightforward. Moreover, it is unclear *a priori* that a sequence-guided drug embedding would translate to a structure-based context.

Surprisingly, we find that they do translate, only requiring a single non-linear projection. Concretely, the pretrained ligand vector 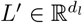 is mapped to a surface-compatible embedding 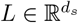 through a learned non-linear transformation, yielding Precise ‘s final processed ligand representation.

### 3.3 Integrating ligand and surface representations to obtain patch scores

The surface and ligand representations 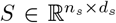 and 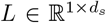 were finally integrated through an inner product operation to produce vertex-associated binding likelihoods *p*:

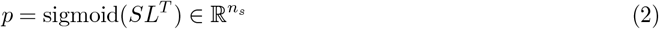

We next aggregate individual vertex scores into patch-level scores, which will be the building blocks of pocket-level analyses. In ligand-docking literature, the minimum patch radius of interest is considered to be 5–9 Å [31]. Choosing the midpoint of that range, we define a patch as a circular region of radius 7 Å centered at some vertex, and compute the patch-level Precise score by averaging per-vertex scores in the patch.

### 3.4 Application 1: binding pocket identification

The patch–ligand prediction function implemented in Precise can be leveraged to identify ligand-specific binding pockets in a protein target. While this is not the primary application we intend for Precise, it *is* valuable for assessing its accuracy as a targeted screening tool. Even for more sophisticated tasks such as docking or protein-ligand co-folding, correctly matching the ligand to its putative pocket is a necessary prerequisite for accuracy in the broader task—correct pose prediction by an ML docking model is irrelevant if the ligand is situated in the incorrect pocket. For pocket prediction with Precise, we could identify the pocket by computing patch scores for all vertices on the surface mesh and aggregating them appropriately. However, this fails to scale. For efficient prediction, we randomly sample 2048 vertices from the surface mesh and compute their corresponding patch scores. Our calculations indicate that 2048 patches are likely to cover the typical protein’s entire surface with a high probability (Supplement A.4). All vertices in a patch inherit its score. If a vertex exists in multiple patches, its score is the average of those patch scores.

Since pockets are typically larger than 7 Å patches, we next aggregate our predictions into a contiguous pocket-like surface region. High binding propensity patches might reside in disjointed but proximal regions. To systematically aggregate such regions into coherent binding regions, we follow LABind [55] and apply the Mean Shift algorithm. It is a non-parametric, mode-seeking clustering method that iteratively shifts data points toward local density maxima and yields distinct, contiguous binding regions. The final binding pocket for the ligand returned by Precise is the centroid of the cluster with the highest average binding score.

### 3.5 Application 2: Precise + Vina for screening of small molecules

To screen a specific target and pocket against a database of compounds, the input to Precise is the target structure and, in that reference frame, the coordinates of the pocket center. We obtain all surface vertices within 4.5Å of center, run the GotenNet surface module on the whole surface and extract the processed surface representations of the vertices of interest. Each vertex is then scored against all 32,768 precomputed discretized drug codes, using our efficient inner product formulation. Per-code scores are averaged across vertices and Precise reports a ranked list of drug codes that are predicted to bind strongly to the target at the pocket. Precise’s modular architecture ensures speed: the geometric deep learning architecture for the surface track is the computationally expensive component, but needs to be computed only once. The ligand encoder can be run quickly across the entire code space with minimal overhead, and the inner product scoring enables screening of 250M compounds in under 2 seconds.

The set of ligands in a high-scoring code comprise candidates for further investigation by docking. However, selecting among these candidates can be a challenge. With large compound databases, each code can map to thousands of ligands (the average Enamine code maps to ∼ 334, 000 ligands [13]). This challenge is not unique to Precise: any uHTVS method likely reports many candidates that need to be selected from.

We introduce an efficient sampling algorithm called Precise-MCTS, which integrates fast Precise-based screening with a selective downstream ‘oracle’ (here, we use Vina-based docking [47,12]) through an iterative Monte Carlo Tree Search (MCTS) approach. Given a protein’s structure and binding site information, Precise-MCTS uses Precise to find the top scoring CoNCISE codes for the site and queries the ZINC 250M database to find a list of potential ligand candidates. These ligands are clustered into an agglomerative tree, with smaller tree distances between ligands with similar binding properties. Finally, a MCMC-like procedure is employed to explore through the tree, zooming in on branches that exhibit strong Vina binding affinities while pruning weak ones, returning an enriched set of ligands that show high Vina-inferred affinity. The complete procedure is detailed in Algorithm 6 (See Supplement).

The key advantage of Precise-MCTS lies in its flexibility: it can efficiently traverse extremely large ligand libraries while giving users granular control over the search trajectory and the evaluation budget (here, the total number of Vina docking runs permitted). The exploration strategy operates on a per-depth basis. At each depth, Precise-MCTS identifies the node with the best Vina docking score (measured on a randomly-chosen descendant leaf), collects all nodes whose scores fall within a range (“slack”) of this best value, and restricts subsequent exploration to this subset. The slack varies with tree depth (Table A.1): Precise employs broader slack near the root to encourage exploration, and progressively tightens it at deeper levels to drive more focused exploitation.

We benchmarked Precise-MCTS against two alternatives. The first is a naive baseline that uses the agglomerative tree solely to partition the candidate space into diverse clusters, returning a random representative from each cluster as the filtered ligand set. The second is Rag2Mol [54], a screening and generation pipeline that begins with a synthesizable drug space and optimizes toward higher-affinity ligand candidates. We evaluated two Precise-MCTS configurations—Precise-MCTS-shallow and Precise-MCTS-deep— where the ‘deep’ setting admits a larger pool of high-scoring codes and ligands, and its per-depth slack allows for a more liberal exploration than the ‘shallow’ setting (this is governed by the ‘TRIM’ function in Algorithm 6, see Supplement A.7 for more detail). As detailed below, the benchmarking results show that Precise-MCTS is a fast, scalable approach that outperforms generative methods even while operating under the more constrained regime of screening within an existing drug space.

### 3.6 Datasets & Training

The model was trained on the PLINDER dataset [11], a well-curated collection of 449,383 protein–ligand interaction (PLI) systems. We note that PLINDER contains holo 3D structures for all the PLI systems in its benchmark, and that we need 3D structure of the targets for training Precise, for input to Precise when testing its performance, and also to construct the ground truth answers to the pocket prediction task to measure its performance. The default train, validation, and test splits provided by PLINDER were used in Precise for model training, hyperparameter optimization, and evaluation (Supplement B.1).

The core Precise model was trained to perform the surface–ligand association prediction task. During training, the input protein–ligand structural coordinates were used to identify the ground-truth surface patches corresponding to ligand binding sites. In addition to these positive examples, random non-binding surface regions were sampled as negative examples. The model was thus trained to distinguish binding patches from non-binding ones. The additional data processing steps and the train, validation and test AUPR scores are reported in Supplementary Figure A.1.

#### Hyperparameters and Training Details

Precise is a compact model, with only **16.9 million** parameters. When the pretrained CoNCISE drug encoder is excluded, the total parameter count reduces further to **11.4 million**. The Precise model employed the default GotenNet hyperparameters for constructing the surface block. The drug encoding block was adopted directly from CoNCISE, with its weights frozen during training. Since the default GotenNet surface representation has a non-steerable dimension of size 256, the input CoNCISE drug embeddings were projected to match this dimensionality, enabling a valid inner product operation. The choice of a 7 Å surface patch was directly inspired by MaSIF [15]. The model was trained for 10 epochs on a single Nvidia H200 GPU using PyTorch’s default AdamW optimizer with a learning rate of 1 × 10^−4^. To account for the imbalance between positive and negative examples we use a focal loss term; following Lin et al.’s recommended defaults [29], we used *α* = 0.7 and *γ* = 2. After 10 epochs (runtime: 30 hours), Precise yielded a validation AUPR of 0.76 (random baseline’s AUPR: 0.091) on the elementary task of surface-ligand binding prediction.

## 4 Results

We first assessed whether Precise could be applied to binding pocket prediction: estimating where a given ligand binds to a given target. We next investigated the accuracy and speed of Precise on the task of screening a large database. Finally, we examined whether a model trained exclusively on monomeric protein surfaces can be applied, in a fully zero-shot manner, to score composite surfaces, including pockets containing metal ions and those formed at multi-chain interfaces.

### 4.1 Application 1: Binding pocket prediction

The binding pocket evaluations were benchmarked against surface-based docking (LABind [55]), generative diffusion-based docking (DiffDock-L [6]), and co-folding approaches (Chai-1 [46], Boltz-2 [39]). We note that the latter three tools address broader tasks than pocket prediction alone: DiffDock-L predicts full ligand poses, while Chai-1 and Boltz-2 predict complete protein–ligand complexes. However, accurate pocket identification is a necessary prerequisite for success in their respective tasks. Our evaluation therefore focuses on pocket prediction as an essential component of effective screening tools. To assess whether ligand quantization impacts pocket prediction performance, we also compared Precise against a variant trained with the full ligand fingerprint representations (termed ‘Precise (no codes)’ in Figure 2) rather than CoNCISE codes.

We evaluated Precise using protein–ligand samples from PLINDER’s evaluation split [11], and further validated results on a newer, independent dataset from RCSB [4] (Supplement B.1). The RCSB dataset, comprising structures deposited after December 2024, provides additional out-of-sample validation, though limited data availability precluded homology-based filtering. Performance was measured as the distance between the predicted and true pocket centers.

Precise either matches or outperforms other methods on the PLINDER test set (Figure 2a). The minimal differences between Precise and ‘Precise (no codes)’ across datasets indicate that ligand discretization via codes does not materially affect accuracy while massively improving scalability. These findings mirror what was observed also in CoNCISE [13]. Achieved here by a model with fewer than 20 million parameters, they underscore the strength of our surface-based representation in capturing target-specific ligand-binding associations.

On the RCSB dataset, both versions of Precise continue to outperform LABind and substantially outperform DiffDock-L; the latter’s performance declines substantially on this dataset. However, Chai-1 and Boltz-2 perform somewhat better on this dataset (Figure 2b). Because Precise is more than 25× faster than Chai-1 and Boltz-2, we recommend applying Precise for initial screening, then filtering top hits with the slower but more comprehensive tools. Again confirming that ligand discretization does not hurt performance, ‘Precise (no codes)’ performs worse than standard Precise.

As a test of Precise’s sensitivity to subtle structural changes, we evaluated whether the learned surface representations could correctly predict shifts in binding affinity induced by single point mutations. For this experiment, we used the PlatinumDB database [41], which provides protein–ligand complex structures paired with experimentally measured binding affinities for both wild-type and mutant variants. We restricted our evaluation to single point mutations for which corresponding PDB structures were directly available from the PlatinumDB website. This yielded a total of 22 complexes; 4 cases had increased and 18 had decreased pocket binding affinities due to mutation. Figures 2(c,d) illustrate two such examples (full results in Supplement B.2) where Precise identified mutational changes in binding fitness that corresponded to experimentally observed results. In Figure 2(c), the human serum albumin pocket shows higher ligand-binding activity in the mutant relative to the wild type (0.30 versus 0.49 in Precise scores). Conversely, Figure 2(d) shows a dihydrofolate reductase pocket where the decrease in Precise score (0.41 for wild type versus 0.38 for the mutant) aligns with the experimentally observed reduction in binding affinity.

Taken together, these findings indicate that Precise can accurately discern ligand-specific pocket preferences and that its pocket-scoring function encodes meaningful information about localized binding fitness, two essential prerequisites for an effective targeted ligand screening pipeline.

### 4.2 Application 2: Pocket-directed ultra high-throughput virtual screening

Effective virtual screening—our primary intended use-case—must balance three competing demands: comprehensive exploration of chemical space, computational feasibility, and practical synthesizability of discovered candidates. We directly evaluated Precise-MCTS (Precise equipped with a Monte Carlo tree search heuristic) as addressing these challenges.

We benchmarked both shallow and deep versions of Precise-MCTS against a naive clustering baseline and Rag2Mol [54] using 65 protein–site pairs from the Rag2Mol study. Rag2Mol employs CoNPLex [44] to screen a 4.5M compound subset of ZINC, performs thousands of docking evaluations, then uses generative optimization to produce synthetic candidates requiring custom synthesis. In contrast, Precise-MCTS screens the ZINC 250M database (50× larger), uses fewer docking evaluations, and returns commercially available compounds. This evaluation setting thus favors Rag2Mol: it operates on a smaller, pre-filtered space with greater computational investment and freedom to generate novel chemistry. Nevertheless, we find that Precise-MCTS outperforms Rag2Mol on Vina docking scores (Figure 3b).

**Fig. 3:**
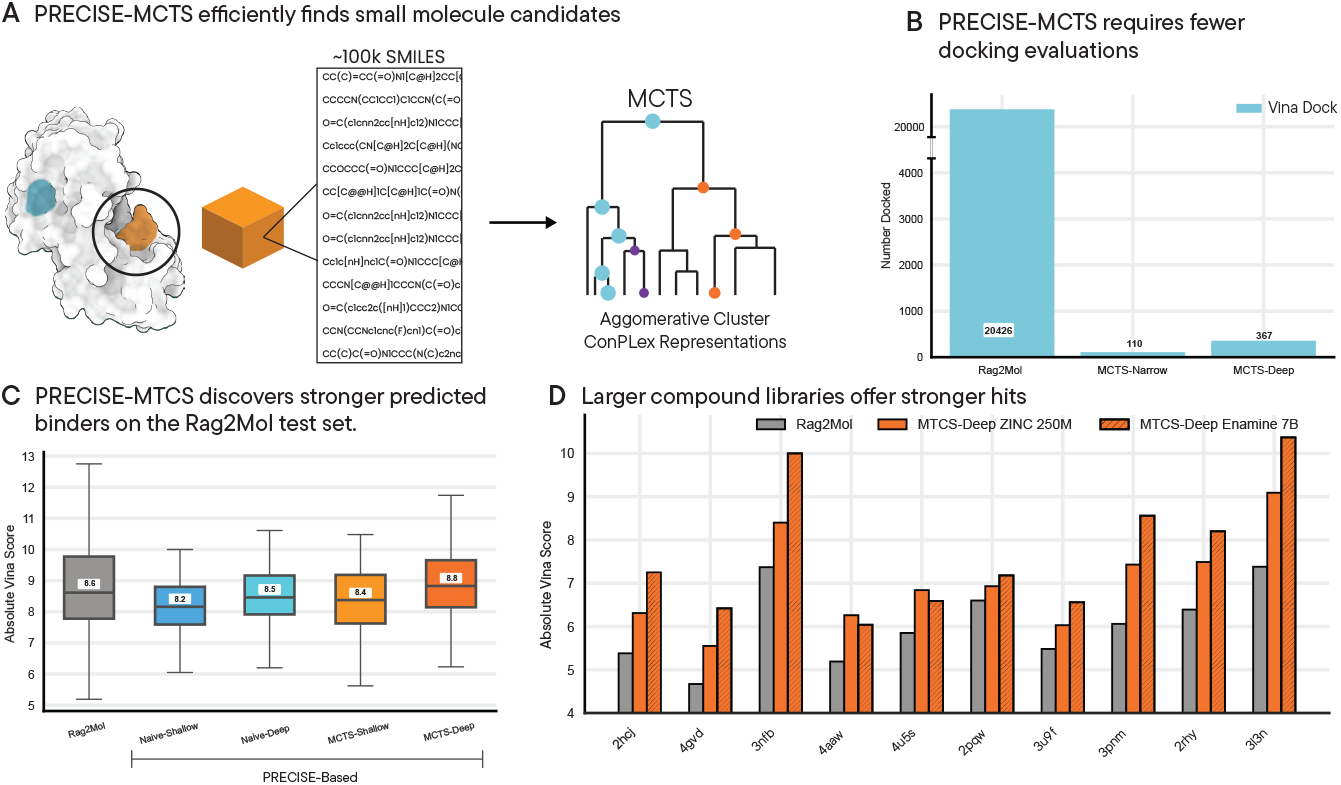
**Precise-MCTS** A) Precise-MCTS uses Precise to identify site-specific codes, retrieves ligands mapped to those codes, and employs an efficient tree-based Monte Carlo sampling pipeline to select small-molecule candidates with high Vina scores. B) Precise-MCTS gives the user control over the number of computationally expensive docking runs during screening; by default, its docking usage is minimal compared to Rag2Mol. C) On the ZINC (250M) database, Precise-MCTS outperforms Rag2Mol on 65 test proteins, even when constrained to provide ligands only from the database. D) For the 10 examples with the weakest Rag2Mol-inferred ligand binders, increasing the database size from ZINC (650M) to Enamine (7B) led to a significant improvement in the best Vina docking scores returned by Precise-MCTS.

For each protein–site pair, each screening approach produced a set of ligand candidates, which we subsequently docked using AutoDock Vina [12]. We report the best (lowest) binding energy for each method, along with the number of docking evaluations required (Figures 3c,d).

We next directly evaluated Precise-MCTS (Precise equipped with a Markov chain tree search heuristic) as a fast and effective drug screening tool—the main use case for which Precise was designed for.

Even the naive selection strategy built on Precise codes performs comparatively well, with MCTS refinement providing substantial additional gains. The naive clustering baseline, which uses Precise to identify high-scoring codes but then performs only simple clustering of ligands in a code and random selection from those candidates, yields ligands with Vina scores competitive with Rag2Mol-generated synthetic compounds (Figure 3). This indicates that the base Precise codes themselves capture high-quality binding-relevant information. Replacing naive selection with the structured Precise-MCTS yields substantial further improvements, outperforming Rag2Mol despite operating under stricter search constraints. Notably, Precise-MCTS achieves these gains while requiring substantially fewer docking evaluations (Figures 3c,d), underscoring both its efficiency and effectiveness.

We further examined whether Precise-MCTS could effectively leverage larger chemical libraries. We replaced ZINC (250M) with Enamine (7B) and re-ran screening on the ten proteins that previously yielded the weakest Rag2Mol Vina scores, maintaining the same docking budget between runs. As shown in Figure 3d, Precise-MCTS achieves higher Vina scores in 8 of 10 cases when using Enamine. Strikingly, the performance gains derive purely from MCTS’s ability to explore the expanded search space: the 25× database increase required only 80% more processing time (median 2h15m for Enamine versus 1h15m for ZINC), with the docking budget held constant.

Together, these results position Precise-MCTS as a powerful and scalable screening framework—one that rivals, and often surpasses, generative approaches while maintaining computational efficiency and delivering immediately synthesizable candidates.

### 4.3 Application 3: Zero-shot generalization to composite targets

Many therapeutically relevant targets exist as composite structures: metalloproteins comprise approximately one-third of all enzymes, protein-protein interfaces present opportunities for fine-grained modulation, and targeting post-translational modifications offers disease specificity. Precise’s surface-based abstraction enables zero-shot generalization to composite targets without model reconfiguration or retraining. We assessed this capability through two experiments on surfaces Precise was never trained on: metal-ion-containing binding sites and multi-chain interfaces.

#### Precise generalizes to metalloprotein surfaces

To quantify Precise’s zero-shot performance on metalion-containing surfaces, we randomly selected 40 proteins from the MetalProGNet training set [22]. These examples feature protein–ligand interactions in which the binding site is activated by a metal ion. For each protein, we computed Precise scores with and without the ion present. The paired comparison (Figure 4b) shows consistently higher scores in the presence of metal ions (*p* = 0.03, one-sided t-test). While modest, this statistically significant effect demonstrates that Precise captures meaningful signal from metal-ion contributions to binding fitness, despite never being trained on such structures.

**Fig. 4:**
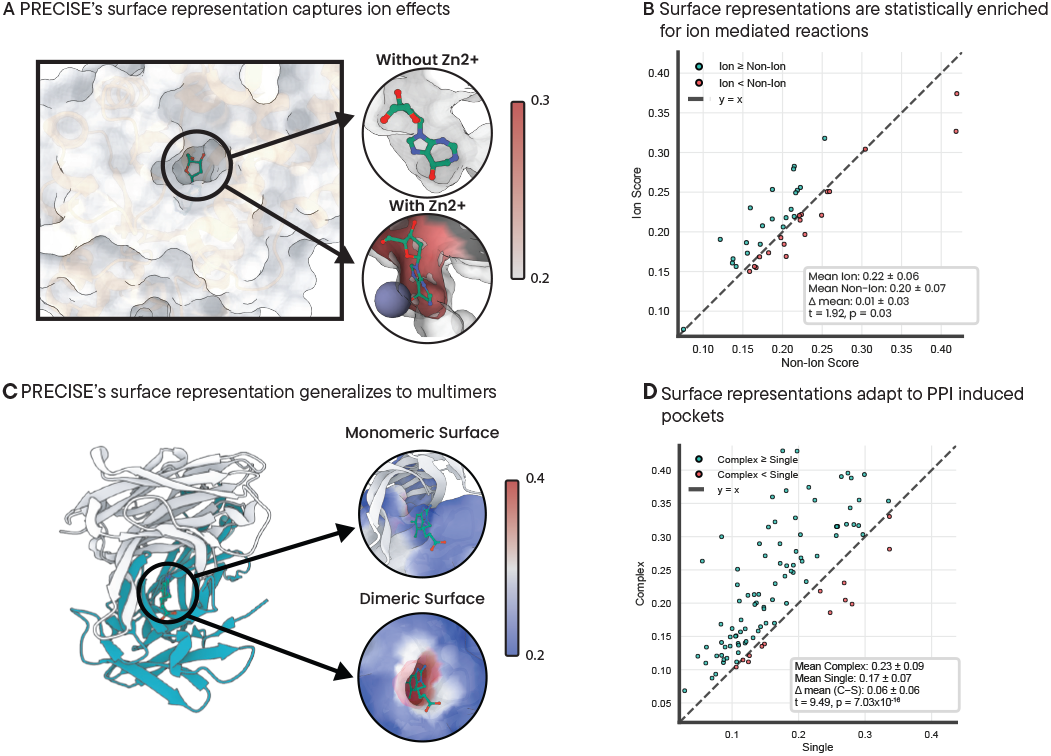
**Surface modifications** A) Precise ‘s surface representation correctly predicts improvements in the ligand affinity of adenosine deaminase (ADA) in a zero-shot manner, when a *Zn*^+2^ ion is introduced near the binding site via MaSIF-NeoSurf. B) Across 40 surface–ion examples randomly sampled from MetalProGNet, Precise binding scores are significantly higher than those of the corresponding non-ionized versions. C) Precise generalizes to multimers and can locate binding spots induced through PPIs. Illustrated are the heavy and light chains of a catalytic elimination antibody (1Y0L) in complex with the small molecule benzimidazolium hapten [9]. This antibody requires both chains to elicit immune response when it comes in contact with the ligand. D) On complexes derived from TernaryDB, Precise ligand-binding scores are enriched for PPI-induced pockets.

As a case study, we examined *Zn*^2+^-mediated ligand binding in adenosine deaminase (ADA) [48]. Using MaSIF-NeoSurf’s surface embeddings [31], we constructed ADA surfaces both with and without the *Zn*^2+^ ion. The presence of *Zn*^2+^ markedly increased the predicted pocket binding affinity (Figure 4a), consistent with the ion’s known role in stabilizing ligand binding.

#### Precise generalizes to multi-chain complex interfaces

We evaluated whether Precise could generalize to multi-chain complexes by benchmarking on TernaryDB [50], which contains structures where ligands bind at shared interfaces between protein chains. For 100 randomly sampled examples, we compared pocket scores on the full multi-chain complex against scores obtained after removing Chain B and treating the structure as monomeric. The results reveal clear and statistically significant divergence between the two settings (Figure 4d). This demonstrates Precise’s ability to capture composite interface features despite training exclusively on monomeric inputs.

As a specific example, Figure 4c shows both heavy and light chains of an antibody (PDB: `1Y0L`) in contact with the small molecule 2-amino-5,6-dimethyl-benzimidazole-1-pentanoic acid (benzimidazolium). At the binding interface, Precise assigns scores of ∼0.4 when both chains are present versus ∼0.2 when only Chain A is considered—a clear preference for the complex form, consistent with experimental evidence that both chains are required for binding.

Together, these results demonstrate that Precise’s zero-shot generalization unlocks previously inaccessible target classes for ultra-high-throughput virtual screening, enabling mechanism-aware searches against metalloproteins, protein-protein interfaces, and other composite structures without requiring target-specific retraining.

## 5 Discussion

Precise addresses a fundamental limitation in ultra-high-throughput virtual screening (uHTVS): existing methods can rank ligand candidates but offer no mechanistic understanding of where molecules bind or how to target specific sites. By operating on protein surface representations enriched with local physicochemical properties, Precise enables pocket-directed screening of billion-compound libraries. This mechanism-aware approach requires the target’s 3D structure as input but, unlike sequence-based binary DTI predictors, can identify binding sites and restrict searches to user-specified pockets.

The surface-based abstraction provides an important benefit: automatic zero-shot generalization to complex targets. Once the surface mesh is characterized, Precise discards all sequence information and operates purely on local geometric and physicochemical properties. A metalloprotein is simply a surface with altered electrostatic patterns; a protein-protein interface presents as extended surface geometry spanning multiple chains. This formulation naturally handles complex targets–—fusion proteins, post-translational modifications, oligomeric assemblies—without model reconfiguration or retraining.

Critically, CoNCISE’s drug codes transfer effectively to this structure-based task despite being learned from sequence-based DTI data. A lightweight linear projection suffices to map codes into the surfacecompatible embedding space. This cross-modal transfer suggests that the hierarchical codes—learned from binary DTI data with only protein sequence inputs—capture fundamental principles of molecular recognition rather than simply sequence artifacts. They thus provide a powerful way to connect two data modalities of DTI studies: experimental binding affinity assays vs those based on structure elucidation. The drug representation transferability enables substantial data efficiency. Pre-learned drug representations mean Precise requires only four surface features per vertex (3 physicochemical, 1 shape-related) to achieve competitive performance. The resulting model contains just 17 million parameters yet provides robust, generalizable accuracy, suggesting these local properties capture the essential physics of molecular recognition when combined with rich drug representations.

Precise-MCTS addresses a critical pain-point in uHTVS: how to prioritize candidates after an initial ranking. The latent drug representations, sourced from CoNPLex [44], enable construction of an agglomerative tree linking structurally-similar molecules. Monte Carlo tree search then efficiently explores this space, using Vina docking scores as an oracle to guide refinement toward high-affinity regions. This framework generalizes beyond Vina: any scoring function (Rosetta energy, MD simulations etc) can serve as the or-acle. The approach relies fundamentally on pre-learned representations that organize chemical space in a binding-relevant manner, enabling efficient traversal without exhaustive enumeration.

The design of Precise implies certain limitations. We work with rigid-body conformations, consistent with most docking methods, and do not model apo-to-holo conformational changes. The limited availability of matched apo-holo training data makes it unclear whether such approaches substantially improve upon rigidbody assumptions. Future work could learn surface representations that predict conformational responses to ligand binding, potentially by training on molecular dynamics trajectories of binding events.

## Supporting information

PRECISE supplement

## 5.1 Code and Data availability

The Precise code is publicly available at https://github.com/rohitsinghlab/PRECISE.

## Acknowledgements

We thank the Tufts BCB group for helpful comments. M.E. was partially supported by ARO 80093-CH-MUR and DARPA grant DOD HR001121S0039

## Disclosure of Interests

M.E., K.D., L.C., and R.S. have filed patents related to this work with provisional patent (#63/735,991).

## A Model and surface description

In this supplementary section, we describe the overall model architecture, including the three main Precise blocks: surface, drug and combined. We further demonstrate how the surface mesh representations were translated into a form suited for a graph-attention based network. Finally, we describe in detail the process by which invariant surface features encoding physico-chemical properties were extracted from the structural data.

### A.1 Precise architecture Surface Block

As summarized in the main paper, the steerable and non-steerable surface features are mainly processed through the GotenNet architecture. The forward algorithm for the surface block is described below:

#### Algorithm 1

Surface Block

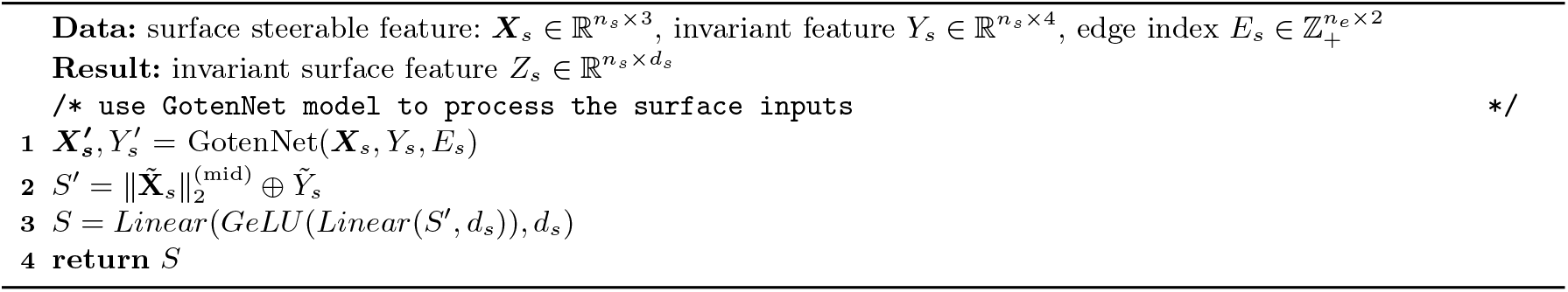

#### How Precise uses GotenNet model

Internally, GotenNet processes the surface features through a graph-attention based, equivariant message passing mechanism. The overall framework begins with the initialization of three parameters derived from the surface inputs: non-steerable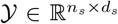, steerable 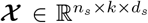 and edge-associated and invariant 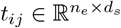 features (*n*_*s*_, *n*_*e*_ denotes the number of surface nodes and edges respectively). Note that, as the default value of *ℓ*_*max*_ (maximum spherical harmonics degree) is set to 1, this leads to the steerable dimension (*k*) being equal to 3. GotenNet initialization is followed by the iterative application of GotenNet’s specialized GATA module (described in more detail below), followed by an equivariant feed-forward network, until the final processed steerable and invariant embeddings are returned as output (Algorithm 2).

##### Algorithm 2

High-level description of GotenNet architecture used in Precise

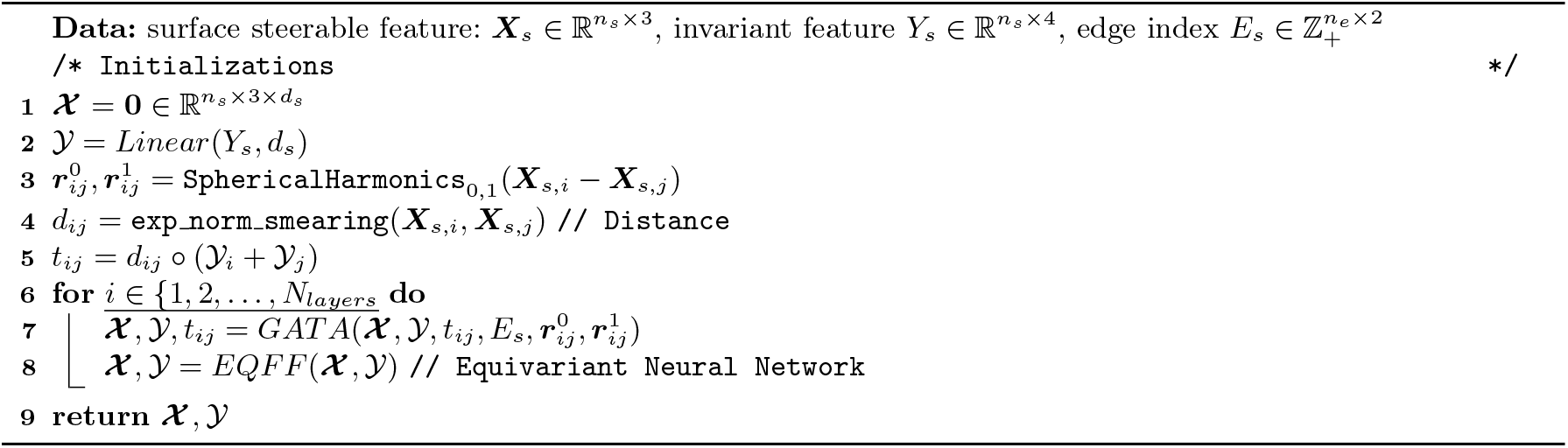

The GATA module in GotenNet does the principle job of message passing and mixing information between the steerable and non-steerable features. The high-level description of the GATA module is described in Algorithm 3. This algorithm is intended as an abstract description of GotenNet, omitting various minor reshaping and transformation steps for clarity. For more implementation details, consult our Precise Github page.

#### Drug encoding block

The simple drug encoder block takes the CoNCISE discretized code embeddings and linearly projects it so that the resulting dimension matches that of the surface representation. This process is described in Algorithm 4

##### Algorithm 3

High level overview of GATA module

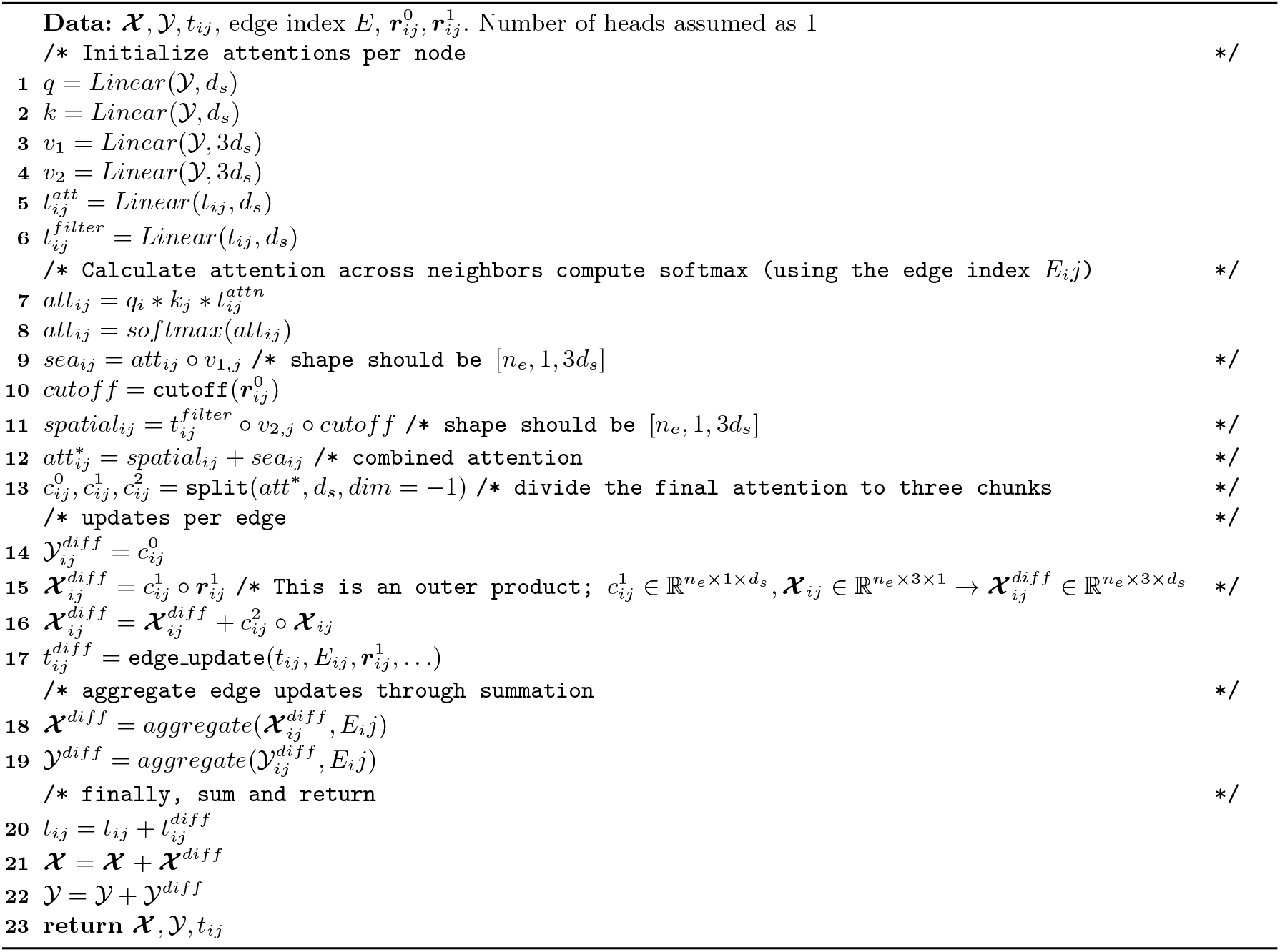

##### Algorithm 4

Precise drug block

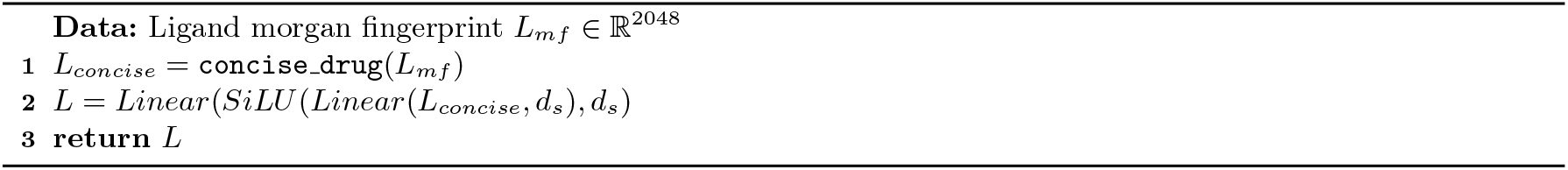

#### The complete Precise architecture

The complete Precise architecture, that takes in a surface representation and ligand Morgan fingerprints and returns a per-vertex surface-ligand binding likelihood is shown below.

##### Algorithm 5

The complete Precise model

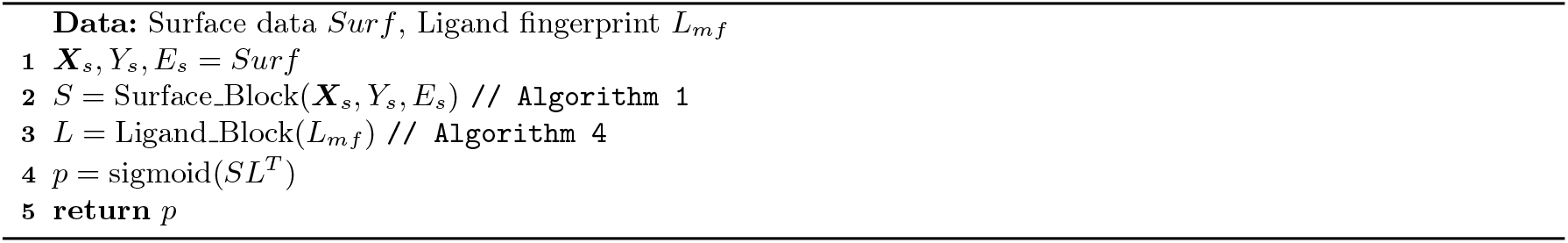

### A.2 Surface Mesh Calculations and Featurizations

The first step in the surface construction pipeline was the addition of hydrogen molecules in a physicochemically stable way using a protonation tool called `Reduce` [49]. Following this, we used `MSMS` [43] to triangulate the molecular surface, applying standard parameters for vertex density (3.0) and water probe radius (1.5 Å). The resulting triangulated surface was then downsampled to a target edge length of approximately 1 Å using the `trimesh` [8] and `pymeshlab` [36] libraries. This processed mesh served as the substrate for calculating four key surface properties: Poisson–Boltzmann continuum electrostatics, proton donor/acceptor characteristics, hydropathy, and shape index.

#### Poisson–Boltzmann continuum electrostatics

The first step in computing local surface electrostatics involved pre-processing the structural file using the `PDB2PQR` tool [10]. Following this, the Poisson-Boltzmann electrostatic potential for each protein molecule was calculated using the `APBS` [3] (v.3.4.1) suite.

The resulting `APBS` output assigned charge information to every vertex on the generated mesh surface, using the `Multivalue` utility within `APBS` for the computation. Finally, the charges were capped between –30 and +30 and then normalized to fall within the range –1 to +1.

#### Free electrons and proton donors

On the `MSMS`-constructed surface, a vertex was classified as a potential donor or acceptor if its closest associated atom was a polar hydrogen, a nitrogen, or an oxygen atom. Based on the orientation of the heavy atoms, a value was then assigned to each vertex, drawn from a Gaussian distribution [26]. This value scale ranged from −1 (indicating the optimal position for a hydrogen bond acceptor) to +1 (indicating the optimal position for a hydrogen bond donor).

#### Hydropathy

Each vertex on the molecular surface also assigned a scalar hydropathy value based on the Kyte and Doolittle scale [27]. The specific value was determined by identifying the amino acid to which the vertex’s closest atom belonged.

The original values from this scale ranged from −4.5 (most hydrophilic) to +4.5 (most hydrophobic). Consistent with the other surface metrics, these hydropathy values were also normalized to a final range of −1 to 1.

#### Shape index

The shape index is a scalar value that describes the local topography of the protein surface at each vertex, with respect to its local curvature [52]. This measure is defined by the principal curvatures, *κ*_1_ and *κ*_2_ (where *κ*_1_ ≥ *κ*_2_).

To calculate this property, we first computed the Mean (*H*) and Gaussian (*K*) curvatures for each vertex on the mesh using the `PyVista` library. From these values, the principal curvatures (*κ*_1_ and *κ*_2_) were derived for each vertex. The final shape index (*SI*) was then computed using the standard formula:

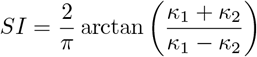

During this computation, a small epsilon (1×10^−8^) was added to the denominator to maintain numerical stability in flat or spherical regions where the principal curvatures may be nearly equal. This calculation directly yields a normalized value ranging from −1 (highly concave shapes) to +1 (highly convex shapes).

### A.3 Calculation of surface metrics on composite surfaces containing ions

The above pipelien was specific to protein-only calculations. However, for complexes containing monatomic ions (e.g., Zn^2+^, Mg^2+^), a modified protocol was implemented. For geometric features (shape index, curvature), the ion’s surface vertices were computed analogously to the protein. For the chemical features (hydropathy, H-bond potential), which are defined by amino acid or organic functional group properties, vertices on the ion surface were assigned default neutral values, as these metrics were not directly applicable. The most significant adaptation occurred in the Poisson–Boltzmann continuum electrostatics pipeline.

A custom script enacted a branching logic where standard (non-metal) organic ligands were parameterized automatically by `PDB2PQR` using the ‘–ligand’ flag. In contrast, for monatomic ions, `PDB2PQR` was executed *without* the ‘–ligand’ flag. Our script then parsed the ion’s MOL2 file, mapped its atom type to a predefined formal charge and PARSE-compatible radius, and appended this information as a new ‘HETATM’ record to the `PDB2PQR` output. This augmented PQR file, integrating both protein and ion, was then used as the input for `APBS`’s `multivalue` tool, ensuring the ion’s electrostatic contribution was accurately incorporated.

### A.4 Number of patches to sample

A good number *n* of vertex induced patches should satisfy the following criteria:

- With high likelihood cover the entire solvent accessible surface area (SASA).
- A random vertex *v* should be represented by multiple patches.

In principle the SASA will increase with the length of the sequence. To find a good choice of *n*, we take the average length of a protein to be approximately 350 amino acids [35]. The average SASA of a protein of this length is around 15000Å^2^ [34].

Assuming points are sampled uniformly on the surface of size *A*, the probability that a vertex *v* is covered by a patch of radius r is given by:

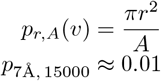

The probability of not covering a vertex with *n* samples is then defined as:

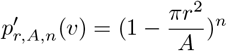

Setting *n* = 2048 under these conditions indicates the probability of **not** being covered is ≈ 1 × 10^−9^.

Now we move onto answering the question: How many patches do I expect *v* to be a part of. Again under the scenario that vertices are selected uniformly, the probability distribution of participating in multiple matches follows the Binomial distribution.

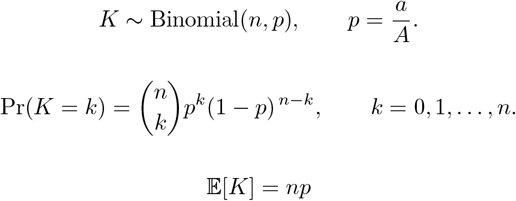

Plugging in for our choices of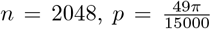, we see that for an average protein ≈ 20 patches participate in the overall include vertex *v*.

### A.5 Training and Validation AUPR results from Precise training

Precise was trained on the PLINDER training dataset, with the validation statistics (AUPR and AU-ROC scores) reported twice every epoch. These metrics are provided in Figure A.1.

**Fig. A.1:**
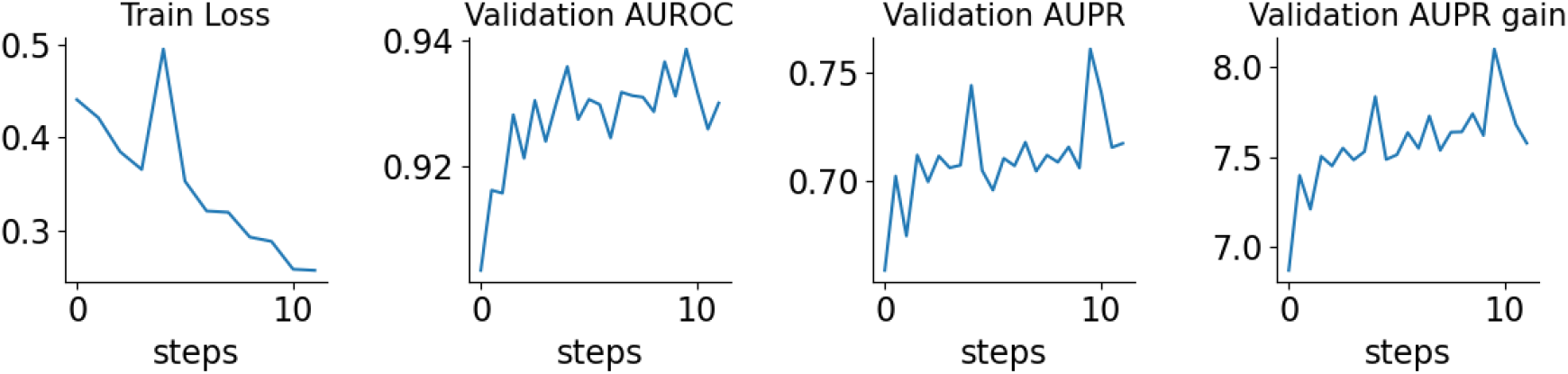
Grid search experiments of Precise on PLINDER evaluation set.

In addition to standard evaluation metrics, we also assessed how well our model performs relative to a randomized baseline on the validation datasets. Specifically, we computed the ratio between the validation AUPR achieved by Precise and that of a theoretical randomized baseline, which we refer to as the AUPR gain. Using this metric as our model-selection criterion, we selected the epoch-10 checkpoint as the default Precise model.

**Fig. A.2:**
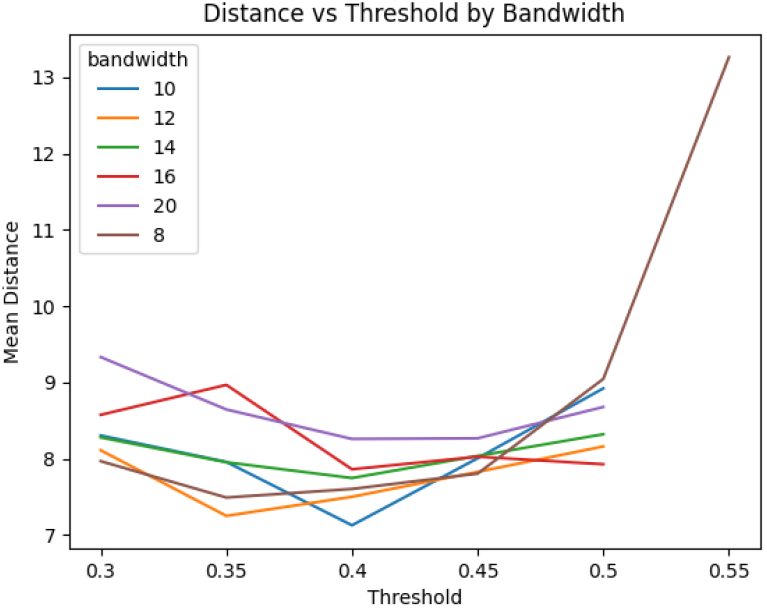
Train and validation statistics obtained from Precise training. For ‘Validation AUPR gain’, we computed the ratio of validation AUPR obtained and the AUPR of a randomized algorithm

### A.6 Ablations of Precise Mean-shift bandwidth and threshold for binding pocket detection

We run ablations on the two key parameters for pocket detection, one being the bandwidth of the meanshift density kernel, the other being the threshold in which Precise’s predictions are considered positive. Although a bandwidth of 10Å at threshold 0.4 reports the lowest mean distance, we choose to take a bandwidth of 12Å at a threshold of 0.35 as it appears to produce a more robust pairing.

#### Algorithm 6

Precise-MCTS

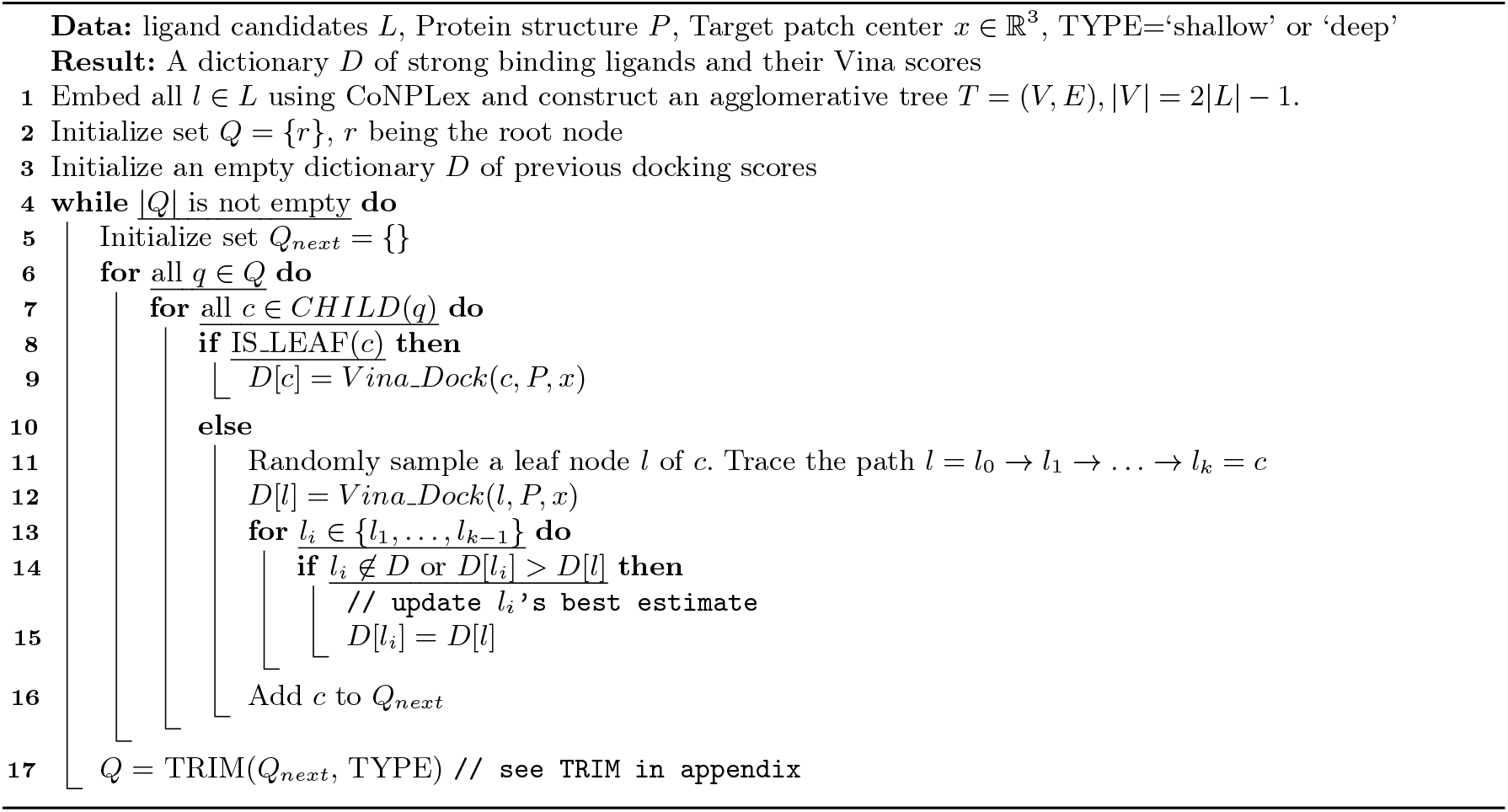

### A.7 Precise-MCTS

### A.8 Construction of naive screening baselines

In addition to Rag2Mol, we compared Precise-MCTS with two naive Precise-based baselines. These baselines were generated by first constructing the agglomerative tree using the ligands’ ConPLex representations (as in Precise-MCTS), then directly clustering the ligands and selecting a representative candidate from each cluster for docking. For both Precise-MCTS and the naive methods, the linkage method was set to ward. Additionally, for the naive deep and shallow clustering methods, the distance thresholds were set to 0.4 and 0.7, respectively.

#### General approach for controlling exploration in the agglomerative tree

The general *TRIM* function, which controls Precise-MCTS exploration utilizes per-depth slack to trim nodes with low-binding scores. A custom *TRIM* function where a slack variable is provided for each depth is described in Algorithm 7.

For the ‘shallow’ and ‘wide’ setting, the per-depth slack setting was set the following way:

##### Algorithm 7

TRIM

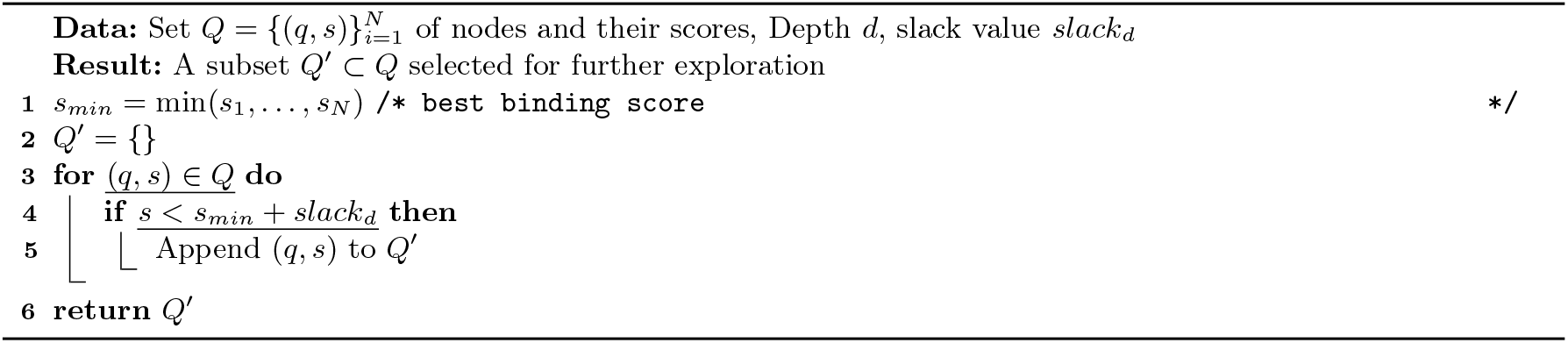

**Table A.1:**
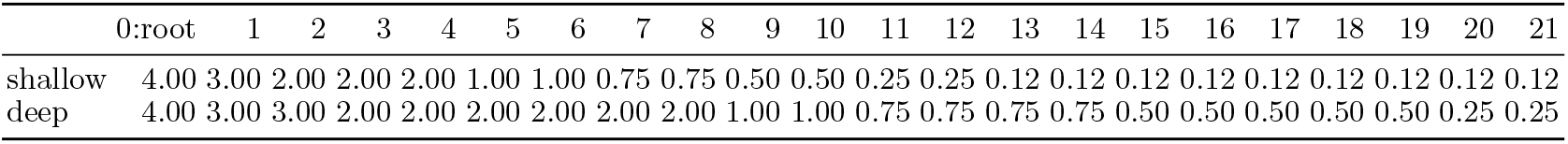
Slack values assigned at each depth 0-21 for the Precise-MCTS shallow and deep settings.

Note that the minimum slack values for the shallow and deep settings are 0.12 and 0.25, respectively. If the search proceeds beyond depth 21, Precise-MCTS continues to use the minimum slack until the search terminates.

#### Additional of additional Precise-MCTS parameters

For both MCTS settings, we selected the top 500 scoring codes. For the shallow setting, we randomly chose 20 ligands from each code to construct the agglomerative tree, whereas for the wide setting, we selected 100 ligands per code. Additionally, in one protein instance where the top 500 codes had minimal ligand coverage (TIAM1 HUMAN 840 931 0 4gvd A), we increased the number of selected codes to 1000 for the deep setting. The full information of the proteins used for the experiment is provided in the GitHub.

## B Dataset description and pre-processing

### B.1 Pocket prediction benchmarking using a newly curated RSCB dataset

We reinforced our pocket prediction results on the PLINDER evaluation dataset by further benchmarking Precise on a newly curated protein–ligand structural dataset sourced from the RCSB PDB. Our primary concern was that some baseline methods might have already been trained on PLINDER PLY examples, potentially giving them an unfair advantage. To mitigate this, we queried the RCSB database to construct a new PLY dataset consisting of monomer–ligand complexes that were released after December 2025, ensuring no overlap with potential training data of baseline models. In addition, we ensured that the queried data were not revisions of structures that were resolved at an earlier date. This returned 253 CIF files in total.

In addition, we queried the CIF files to find the exact name of ligand bound with the proteins. After filtering out candidates with very small ligands (i.e. those with molecular weight *<* 100), we finally obtained 100 PLY structures, which we used as the second dataset for pocket prediction. The PDB id of these 100 structures are provided below:

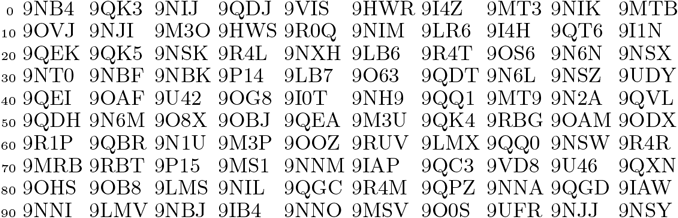

### B.2 Evaluating mutational effects on pocket fitness through PlatinumDB

PlatinumDB, which contains experimentally measured changes in binding affinities wild-type and mutational variants, was used to assess Precise’s sensitivity to surface modifications.

To get the evaluation dataset for the mutational effect, we get all the available data from PlatinumDB [41], which is a structural database of experimentally measured effects of mutations on protein-ligand complexes. The protein surface calculation pipeline (Section 3.1) was applied to each mutation protein as well as the corresponding wild type. Only single amino acid mutations were adopted during the evaluation.

### B.3 PPI Induced Pocket

The TernaryDB dataset [50], which contains ternary structures where the ligand is bound to common interfaces, was used to assess Precise’s zero-shot generalization to multi-chain surfaces. In total, TernaryDB has 22,303 such complexes, out of which we randomly sampled 100 for evaluation. Fitness assessments were done by comparing Precise’s scores on the ligand binding site in the multi-chain interface against scores at the same site after discarding ‘Chain B’. The CSV file describing the 100 TernaryDB examples used for multi-chain evaluations are provided in GitHub.

## Footnotes

4 For computational efficiency, we sample vertices rather than exhaustively scoring the entire surface mesh (Section A.4).

## Notes

### Competing Interest Statement

The authors have declared no competing interest.

