## Supplementary material for "Learning a PRECISE language for small-molecule binding": PRECISE supplement

### A Model and surface description

In this supplementary section, we describe the overall model architecture, including the three main PRECISE blocks: surface, drug and combined. We further demonstrate how the surface mesh representations were translated into a form suited for a graph-attention based network. Finally, we describe in detail the process by which invariant surface features encoding physico-chemical properties were extracted from the structural data.

#### A.1 Precise architecture

##### Surface Block

As summarized in the main paper, the steerable and non-steerable surface features are mainly processed through the GotenNet architecture. The forward algorithm for the surface block is described below:

**Algorithm 1:** Surface Block

**Data:** surface steerable feature: $\boldsymbol{X}_s \in \mathbb{R}^{n_s \times 3}$, invariant feature $Y_s \in \mathbb{R}^{n_s \times 4}$, edge index $E_s \in \mathbb{Z}_+^{n_e \times 2}$
**Result:** invariant surface feature $Z_s \in \mathbb{R}^{n_s \times d_s}$
`/* use GotenNet model to process the surface inputs */`
$\boldsymbol{X}'_s, Y'_s = \text{GotenNet}(\boldsymbol{X}_s, Y_s, E_s)$
$S' = \|\tilde{\mathbf{X}}_s\|_2^{(\text{mid})} \oplus \tilde{Y}_s$
$S = Linear(GeLU(Linear(S', d_s)), d_s)$
**return** $S$

##### How Precise uses GotenNet model

Internally, GotenNet processes the surface features through a graph-attention based, equivariant message passing mechanism. The overall framework begins with the initialization of three parameters derived from the surface inputs: non-steerable $\mathcal{Y} \in \mathbb{R}^{n_s \times d_s}$, steerable $\boldsymbol{\mathcal{X}} \in \mathbb{R}^{n_s \times k \times d_s}$ and edge-associated and invariant $t_{ij} \in \mathbb{R}^{n_e \times d_s}$ features ($n_s, n_e$ denotes the number of surface nodes and edges respectively). Note that, as the default value of $\ell_{max}$ (maximum spherical harmonics degree) is set to 1, this leads to the steerable dimension ($k$) being equal to 3. GotenNet initialization is followed by the iterative application of GotenNet's specialized GATA module (described in more detail below), followed by an equivariant feed-forward network, until the final processed steerable and invariant embeddings are returned as output (Algorithm 2).

**Algorithm 2:** High-level description of GotenNet architecture used in PRECISE

**Data:** surface steerable feature: $\boldsymbol{X}_s \in \mathbb{R}^{n_s \times 3}$, invariant feature $Y_s \in \mathbb{R}^{n_s \times 4}$, edge index $E_s \in \mathbb{Z}_+^{n_e \times 2}$
`/* Initializations */`
$\boldsymbol{\mathcal{X}} = \mathbf{0} \in \mathbb{R}^{n_s \times 3 \times d_s}$
$\mathcal{Y} = Linear(Y_s, d_s)$
$\boldsymbol{r}_{ij}^0, \boldsymbol{r}_{ij}^1 = \texttt{SphericalHarmonics}_{0,1}(\boldsymbol{X}_{s,i} - \boldsymbol{X}_{s,j})$
$d_{ij} = \texttt{exp\_norm\_smearing}(\boldsymbol{X}_{s,i}, \boldsymbol{X}_{s,j})$ `// Distance`
$t_{ij} = d_{ij} \circ (\mathcal{Y}_i + \mathcal{Y}_j)$
**for** $i \in \{1, 2, \ldots, N_{layers}\}$ **do**
  $\boldsymbol{\mathcal{X}}, \mathcal{Y}, t_{ij} = GATA(\boldsymbol{\mathcal{X}}, \mathcal{Y}, t_{ij}, E_s, \boldsymbol{r}_{ij}^0, \boldsymbol{r}_{ij}^1)$
  $\boldsymbol{\mathcal{X}}, \mathcal{Y} = EQFF(\boldsymbol{\mathcal{X}}, \mathcal{Y})$ `// Equivariant Neural Network`
**return** $\boldsymbol{\mathcal{X}}, \mathcal{Y}$

The GATA module in GotenNet does the principle job of message passing and mixing information between the steerable and non-steerable features. The high-level description of the GATA module is described in Algorithm 3. This algorithm is intended as an abstract description of GotenNet, omitting various minor reshaping and transformation steps for clarity. For more implementation details, consult our PRECISE Github page.

##### Drug encoding block

The simple drug encoder block takes the CoNCISE discretized code embeddings and linearly projects it so that the resulting dimension matches that of the surface representation. This process is described in Algorithm 4

**Algorithm 3:** High level overview of GATA module

**Data:** $\boldsymbol{\mathcal{X}}, \mathcal{Y}, t_{ij}$, edge index $E$, $\boldsymbol{r}_{ij}^0, \boldsymbol{r}_{ij}^1$. Number of heads assumed as 1

`/* Initialize attentions per node */`

$q = Linear(\mathcal{Y}, d_s)$

$k = Linear(\mathcal{Y}, d_s)$

$v_1 = Linear(\mathcal{Y}, 3d_s)$

$v_2 = Linear(\mathcal{Y}, 3d_s)$

$t_{ij}^{att} = Linear(t_{ij}, d_s)$

$t_{ij}^{filter} = Linear(t_{ij}, d_s)$

`/* Calculate attention across neighbors compute softmax (using the edge index` $E_ij$`) */`

$att_{ij} = q_i * k_j * t_{ij}^{attn}$

$att_{ij} = softmax(att_{ij})$

$sea_{ij} = att_{ij} \circ v_{1,j}$ `/* shape should be` $[n_e, 1, 3d_s]$ `*/`

$cutoff = \texttt{cutoff}(\boldsymbol{r}_{ij}^0)$

$spatial_{ij} = t_{ij}^{filter} \circ v_{2,j} \circ cutoff$ `/* shape should be` $[n_e, 1, 3d_s]$ `*/`

$att_{ij}^* = spatial_{ij} + sea_{ij}$ `/* combined attention */`

$c_{ij}^0, c_{ij}^1, c_{ij}^2 = \texttt{split}(att^*, d_s, dim = -1)$ `/* divide the final attention to three chunks */`

`/* updates per edge */`

$\mathcal{Y}_{ij}^{diff} = c_{ij}^0$

$\boldsymbol{\mathcal{X}}_{ij}^{diff} = c_{ij}^1 \circ \boldsymbol{r}_{ij}^1$ `/* This is an outer product;` $c_{ij}^1 \in \mathbb{R}^{n_e \times 1 \times d_s}, \boldsymbol{\mathcal{X}}_{ij} \in \mathbb{R}^{n_e \times 3 \times 1} \rightarrow \boldsymbol{\mathcal{X}}_{ij}^{diff} \in \mathbb{R}^{n_e \times 3 \times d_s}$ `*/`

$\boldsymbol{\mathcal{X}}_{ij}^{diff} = \boldsymbol{\mathcal{X}}_{ij}^{diff} + c_{ij}^2 \circ \boldsymbol{\mathcal{X}}_{ij}$

$t_{ij}^{diff} = \texttt{edge\_update}(t_{ij}, E_{ij}, \boldsymbol{r}_{ij}^1, \ldots)$

`/* aggregate edge updates through summation */`

$\boldsymbol{\mathcal{X}}^{diff} = aggregate(\boldsymbol{\mathcal{X}}_{ij}^{diff}, E_ij)$

$\mathcal{Y}^{diff} = aggregate(\mathcal{Y}_{ij}^{diff}, E_ij)$

`/* finally, sum and return */`

$t_{ij} = t_{ij} + t_{ij}^{diff}$

$\boldsymbol{\mathcal{X}} = \boldsymbol{\mathcal{X}} + \boldsymbol{\mathcal{X}}^{diff}$

$\mathcal{Y} = \mathcal{Y} + \mathcal{Y}^{diff}$

**return** $\boldsymbol{\mathcal{X}}, \mathcal{Y}, t_{ij}$

**Algorithm 4:** PRECISE drug block

**Data:** Ligand morgan fingerprint $L_{mf} \in \mathbb{R}^{2048}$

$L_{concise} = \texttt{concise\_drug}(L_{mf})$

$L = Linear(SiLU(Linear(L_{concise}, d_s), d_s)$

**return** $L$

##### The complete Precise architecture

The complete PRECISE architecture, that takes in a surface representation and ligand Morgan fingerprints and returns a per-vertex surface-ligand binding likelihood is shown below.

**Algorithm 5:** The complete PRECISE model

**Data:** Surface data $Surf$, Ligand fingerprint $L_{mf}$

$\boldsymbol{X}_s, Y_s, E_s = Surf$

$S = \text{Surface\_Block}(\boldsymbol{X}_s, Y_s, E_s)$ `// Algorithm 1`

$L = \text{Ligand\_Block}(L_{mf})$ `// Algorithm 4`

$$SI = \frac{2}{\pi} \arctan\left(\frac{\kappa_1 + \kappa_2}{\kappa_1 - \kappa_2}\right)$$

During this computation, a small epsilon ($1 \times 10^{-8}$) was added to the denominator to maintain numerical stability in flat or spherical regions where the principal curvatures may be nearly equal. This calculation directly yields a normalized value ranging from -1 (highly concave shapes) to +1 (highly convex shapes).

##### A.3 Calculation of surface metrics on composite surfaces containing ions

The above pipelien was specific to protein-only calculations. However, for complexes containing monatomic ions (e.g., $Zn^{2+}$, $Mg^{2+}$), a modified protocol was implemented. For geometric features (shape index, curvature), the ion's surface vertices were computed analogously to the protein. For the chemical features (hydropathy, H-bond potential), which are defined by amino acid or organic functional group properties, vertices on the ion surface were assigned default neutral values, as these metrics were not directly applicable.

The most significant adaptation occurred in the Poisson–Boltzmann continuum electrostatics pipeline. A custom script enacted a branching logic where standard (non-metal) organic ligands were parameterized automatically by `PDB2PQR` using the '–ligand' flag. In contrast, for monatomic ions, `PDB2PQR` was executed *without* the '–ligand' flag. Our script then parsed the ion's MOL2 file, mapped its atom type to a predefined formal charge and PARSE-compatible radius, and appended this information as a new 'HETATM' record to the `PDB2PQR` output. This augmented PQR file, integrating both protein and ion, was then used as the input for `APBS`'s `multivalue` tool, ensuring the ion's electrostatic contribution was accurately incorporated.

##### A.4 Number of patches to sample

A good number $n$ of vertex induced patches should satisfy the following criteria:

1. With high likelihood cover the entire solvent accessible surface area (SASA).
2. A random vertex $v$ should be represented by multiple patches.

In principle the SASA will increase with the length of the sequence. To find a good choice of $n$, we take the average length of a protein to be approximately 350 amino acids [35]. The average SASA of a protein of this length is around $15000 \mathring{A}^2$ [34].

Assuming points are sampled uniformly on the surface of size $A$, the probability that a vertex $v$ is covered by a patch of radius r is given by:

$$p_{r,A}(v) = \frac{\pi r^2}{A}$$
$$p_{7\text{Å},\,15000} \approx 0.01$$

The probability of not covering a vertex with $n$ samples is then defined as:

$$p'_{r,A,n}(v) = (1 - \frac{\pi r^2}{A})^n$$

Setting $n = 2048$ under these conditions indicates the probability of **not** being covered is $\approx 1 \times 10^{-9}$.

Now we move onto answering the question: How many patches do I expect $v$ to be a part of. Again under the scenario that vertices are selected uniformly, the probability distribution of participating in multiple matches follows the Binomial distribution.

$$K \sim \text{Binomial}(n, p), \qquad p = \frac{a}{A}.$$

$$\Pr(K = k) = \binom{n}{k} p^k (1 - p)^{n-k}, \qquad k = 0, 1, \ldots, n.$$

$$\mathbb{E}[K] = np$$

Plugging in for our choices of $n = 2048$, $p = \frac{49\pi}{15000}$, we see that for an average protein $\approx 20$ patches participate in the overall include vertex $v$.

##### A.5 Training and Validation AUPR results from Precise training

PRECISE was trained on the PLINDER training dataset, with the validation statistics (AUPR and AUROC scores) reported twice every epoch. These metrics are provided in Figure A.1.

In addition to standard evaluation metrics, we also assessed how well our model performs relative to a randomized baseline on the validation datasets. Specifically, we computed the ratio between the validation AUPR achieved by PRECISE and that of a theoretical randomized baseline, which we refer to as the AUPR gain. Using this metric as our model-selection criterion, we selected the epoch-10 checkpoint as the default PRECISE model.

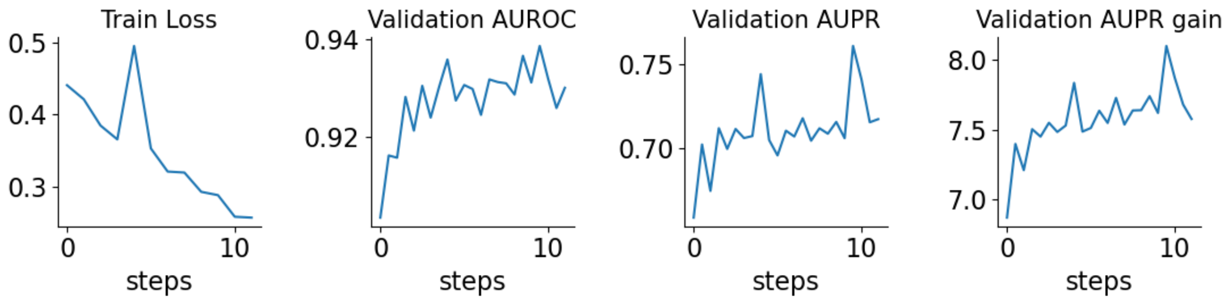


Fig. A.1: Train and validation statistics obtained from PRECISE training. For 'Validation AUPR gain', we computed the ratio of validation AUPR obtained and the AUPR of a randomized algorithm

##### A.6 Ablations of Precise Mean-shift bandwidth and threshold for binding pocket detection

We run ablations on the two key parameters for pocket detection, one being the bandwidth of the mean-shift density kernel, the other being the threshold in which PRECISE's predictions are considered positive. Although a bandwidth of 10Å at threshold 0.4 reports the lowest mean distance, we choose to take a bandwidth of 12Å at a threshold of 0.35 as it appears to produce a more robust pairing.

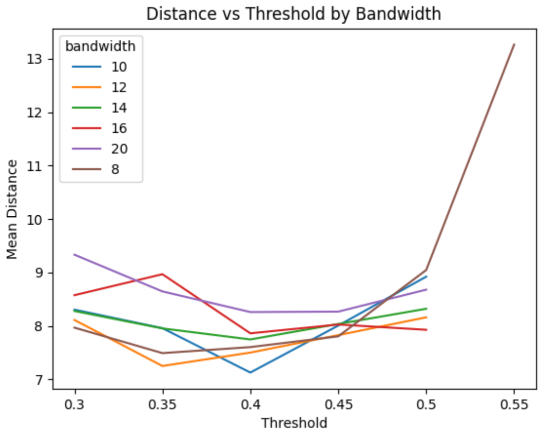


Fig. A.2: Grid search experiments of PRECISE on PLINDER evaluation set.

**Algorithm 6:** PRECISE-MCTS

**Data:** ligand candidates $L$, Protein structure $P$, Target patch center $x \in \mathbb{R}^3$, TYPE='shallow' or 'deep'
**Result:** A dictionary $D$ of strong binding ligands and their Vina scores

Embed all $l \in L$ using CoNPLex and construct an agglomerative tree $T = (V, E), |V| = 2|L| - 1$.
Initialize set $Q = \{r\}$, $r$ being the root node
Initialize an empty dictionary $D$ of previous docking scores
**while** $|Q|$ is not empty **do**
  Initialize set $Q_{next} = \{\}$
  **for** all $q \in Q$ **do**
    **for** all $c \in CHILD(q)$ **do**
      **if** IS_LEAF($c$) **then**
        $D[c] = Vina\_Dock(c, P, x)$
      **else**
        Randomly sample a leaf node $l$ of $c$. Trace the path $l = l_0 \to l_1 \to \ldots \to l_k = c$
        $D[l] = Vina\_Dock(l, P, x)$
        **for** $l_i \in \{l_1, \ldots, l_{k-1}\}$ **do**
          **if** $l_i \notin D$ or $D[l_i] > D[l]$ **then**
            `// update` $l_i$`'s best estimate`
            $D[l_i] = D[l]$
        Add $c$ to $Q_{next}$
  $Q$ = TRIM($Q_{next}$, TYPE) `// see TRIM in appendix`

#### A.7 Precise-MCTS

#### A.8 Construction of naive screening baselines

In addition to Rag2Mol, we compared PRECISE-MCTS with two naive PRECISE-based baselines. These baselines were generated by first constructing the agglomerative tree using the ligands' ConPLex representations (as in PRECISE-MCTS), then directly clustering the ligands and selecting a representative candidate from each cluster for docking. For both PRECISE-MCTS and the naive methods, the linkage method was set to ward. Additionally, for the naive deep and shallow clustering methods, the distance thresholds were set to 0.4 and 0.7, respectively.

##### General approach for controlling exploration in the agglomerative tree

The general $TRIM$ function, which controls PRECISE-MCTS exploration utilizes per-depth slack to trim nodes with low-binding scores. A custom $TRIM$ function where a slack variable is provided for each depth is described in Algorithm 7.

For the 'shallow' and 'wide' setting, the per-depth slack setting was set the following way:

---

**Algorithm 7:** TRIM

**Data:** Set $Q = \{(q,s)\}_{i=1}^{N}$ of nodes and their scores, Depth $d$, slack value $slack_d$
**Result:** A subset $Q' \subset Q$ selected for further exploration

```
s_min = min(s_1, ..., s_N) /* best binding score */
Q' = {}
for (q, s) ∈ Q do
    if s < s_min + slack_d then
        Append (q, s) to Q'
return Q'
```

---

Table A.1: Slack values assigned at each depth 0-21 for the PRECISE-MCTS shallow and deep settings.

| | 0:root | 1 | 2 | 3 | 4 | 5 | 6 | 7 | 8 | 9 | 10 | 11 | 12 | 13 | 14 | 15 | 16 | 17 | 18 | 19 | 20 | 21 |
|---|---|---|---|---|---|---|---|---|---|---|---|---|---|---|---|---|---|---|---|---|---|---|
| shallow | 4.00 | 3.00 | 2.00 | 2.00 | 2.00 | 1.00 | 1.00 | 0.75 | 0.75 | 0.50 | 0.50 | 0.25 | 0.25 | 0.12 | 0.12 | 0.12 | 0.12 | 0.12 | 0.12 | 0.12 | 0.12 | 0.12 |
| deep | 4.00 | 3.00 | 3.00 | 2.00 | 2.00 | 2.00 | 2.00 | 2.00 | 2.00 | 1.00 | 1.00 | 0.75 | 0.75 | 0.75 | 0.75 | 0.50 | 0.50 | 0.50 | 0.50 | 0.50 | 0.25 | 0.25 |

Note that the minimum slack values for the shallow and deep settings are 0.12 and 0.25, respectively. If the search proceeds beyond depth 21, PRECISE-MCTS continues to use the minimum slack until the search terminates.

**Additional of additional Precise-MCTS parameters**

For both MCTS settings, we selected the top 500 scoring codes. For the shallow setting, we randomly chose 20 ligands from each code to construct the agglomerative tree, whereas for the wide setting, we selected 100 ligands per code. Additionally, in one protein instance where the top 500 codes had minimal ligand coverage (TIAM1_HUMAN_840_931_0_4gvd_A), we increased the number of selected codes to 1000 for the deep setting. The full information of the proteins used for the experiment is provided in the GitHub.

### B Dataset description and pre-processing

#### B.1 Pocket prediction benchmarking using a newly curated RSCB dataset

We reinforced our pocket prediction results on the PLINDER evaluation dataset by further benchmarking PRECISE on a newly curated protein–ligand structural dataset sourced from the RCSB PDB. Our primary concern was that some baseline methods might have already been trained on PLINDER PLY examples, potentially giving them an unfair advantage. To mitigate this, we queried the RCSB database to construct a new PLY dataset consisting of monomer–ligand complexes that were released after December 2025, ensuring no overlap with potential training data of baseline models. In addition, we ensured that the queried data were not revisions of structures that were resolved at an earlier date. This returned 253 CIF files in total.

In addition, we queried the CIF files to find the exact name of ligand bound with the proteins. After filtering out candidates with very small ligands (i.e. those with molecular weight $< 100$), we finally obtained 100 PLY structures, which we used as the second dataset for pocket prediction. The PDB id of these 100 structures are provided below:

| | | | | | | | | | | |
|---|---|---|---|---|---|---|---|---|---|---|
| 0 | 9NB4 | 9QK3 | 9NIJ | 9QDJ | 9VIS | 9HWR | 9I4Z | 9MT3 | 9NIK | 9MTB |
| 10 | 9OVJ | 9NJI | 9M3O | 9HWS | 9R0Q | 9NIM | 9LR6 | 9I4H | 9QT6 | 9I1N |
| 20 | 9QEK | 9QK5 | 9NSK | 9R4L | 9NXH | 9LB6 | 9R4T | 9OS6 | 9N6N | 9NSX |
| 30 | 9NT0 | 9NBF | 9NBK | 9P14 | 9LB7 | 9O63 | 9QDT | 9N6L | 9NSZ | 9UDY |
| 40 | 9QEI | 9OAF | 9U42 | 9OG8 | 9I0T | 9NH9 | 9QQ1 | 9MT9 | 9N2A | 9QVL |
| 50 | 9QDH | 9N6M | 9O8X | 9OBJ | 9QEA | 9M3U | 9QK4 | 9RBG | 9OAM | 9ODX |
| 60 | 9R1P | 9QBR | 9N1U | 9M3P | 9OOZ | 9RUV | 9LMX | 9QQ0 | 9NSW | 9R4R |
| 70 | 9MRB | 9RBT | 9P15 | 9MS1 | 9NNM | 9IAP | 9QC3 | 9VD8 | 9U46 | 9QXN |
| 80 | 9OHS | 9OB8 | 9LMS | 9NIL | 9QGC | 9R4M | 9QPZ | 9NNA | 9QGD | 9IAW |
| 90 | 9NNI | 9LMV | 9NBJ | 9IB4 | 9NNO | 9MSV | 9O0S | 9UFR | 9NJJ | 9NSY |

#### B.2 Evaluating mutational effects on pocket fitness through PlatinumDB

PlatinumDB, which contains experimentally measured changes in binding affinities wild-type and mutational variants, was used to assess PRECISE's sensitivity to surface modifications.

To get the evaluation dataset for the mutational effect, we get all the available data from PlatinumDB [41], which is a structural database of experimentally measured effects of mutations on protein-ligand complexes.

The protein surface calculation pipeline (Section 3.1) was applied to each mutation protein as well as the corresponding wild type. Only single amino acid mutations were adopted during the evaluation.

##### B.3 PPI Induced Pocket

The TernaryDB dataset [50], which contains ternary structures where the ligand is bound to common interfaces, was used to assess PRECISE's zero-shot generalization to multi-chain surfaces. In total, TernaryDB has 22,303 such complexes, out of which we randomly sampled 100 for evaluation. Fitness assessments were done by comparing PRECISE's scores on the ligand binding site in the multi-chain interface against scores at the same site after discarding 'Chain B'. The CSV file describing the 100 TernaryDB examples used for multi-chain evaluations are provided in GitHub.
